# Cortico-cerebellar effective connectivity during adapting to vs ignoring delayed visual movement feedback

**DOI:** 10.1101/2025.10.23.684115

**Authors:** Zhenyu Wang, Jakub Limanowski

**Author notes:** **Correspondence:** Jakub Limanowski, Universität Greifswald, Institut für Psychologie, Franz-Mehring-Straße 47, 17489, Greifswald, Germany.

## Abstract

We modelled hemodynamic responses acquired during a virtual reality based hand-target matching task with conditions in which delayed visual movement feedback was behaviorally relevant (requiring visuomotor adaptation) vs irrelevant (i.e., needed to be ignored). We had observed increased hemodynamic responses in the cerebellum, V5, and intraparietal sulcus linked to delay-dependent adaptation. Here, we used dynamic causal modeling to test if these regional activity changes could be explained in terms of network interactions among those nodes. We found a strong excitatory influence of the right cerebellum on the bilateral V5, which increased during the visuomotor adaptation > no adaptation tasks. Furthermore, there was an increased mutual excitation among the cerebellar hemispheres, and an inhibition of the cerebella by the V5, during visuomotor adaptation. These results are consistent with the idea that the cerebellum implements forward models predicting the sensory consequences of actions, and communicates these predictions to other regions of the visuomotor control hierarchy.

## Introduction

On-line motor control relies on flexible internal models in the brain, which, among other things, must entail accurate estimates of the state of the body, and accurate predictions of the sensory consequences that actions will produce (Desmurget & Grafton, 2000; Wolpert & Flanagan, 2001; Shadmehr & Krakauer, 2008). Neurophysiological, brain imaging, and clinical studies suggest that such state estimation and “forward models” issuing sensory predictions may be implemented by the cerebellum and the posterior parietal cortex (PPC, Blakemore et al., 1998; Desmurget et al., 1999; Miall et al., 1993; Wolpert et al., 1998a,b; Synofzik et al., 2008; Izawa et al., 2012). Besides changes in regional (e.g., cerebellar) brain activity, several studies have furthermore demonstrated changes in cortico-cerebellar connectivity during the prediction of sensory action consequences (Kilteni & Ehrsson, 2020; Kellermann et al., 2012; Arikan et al., 2019; Tzvi et al., 2020).

We (Vigh & Limanowski, 2025) recently reported increased PPC, occipitotemporal, and cerebellar BOLD in a task where participants had to adapt their hand movements to repeatedly varying visuomotor delays (i.e., temporal lag added to the visual movement feedback), compared with when they moved under identical visuomotor delays but had to ignore them (Fig. 1). Thereby the cerebellar BOLD was furthermore correlated with the amount of visuomotor control only in the adaptation task; i.e., when visual feedback delays were behaviorally relevant. These results suggested brain areas specifically involved in predicting and processing visual movement feedback depending on its behavioral relevance.

**Figure 1:**
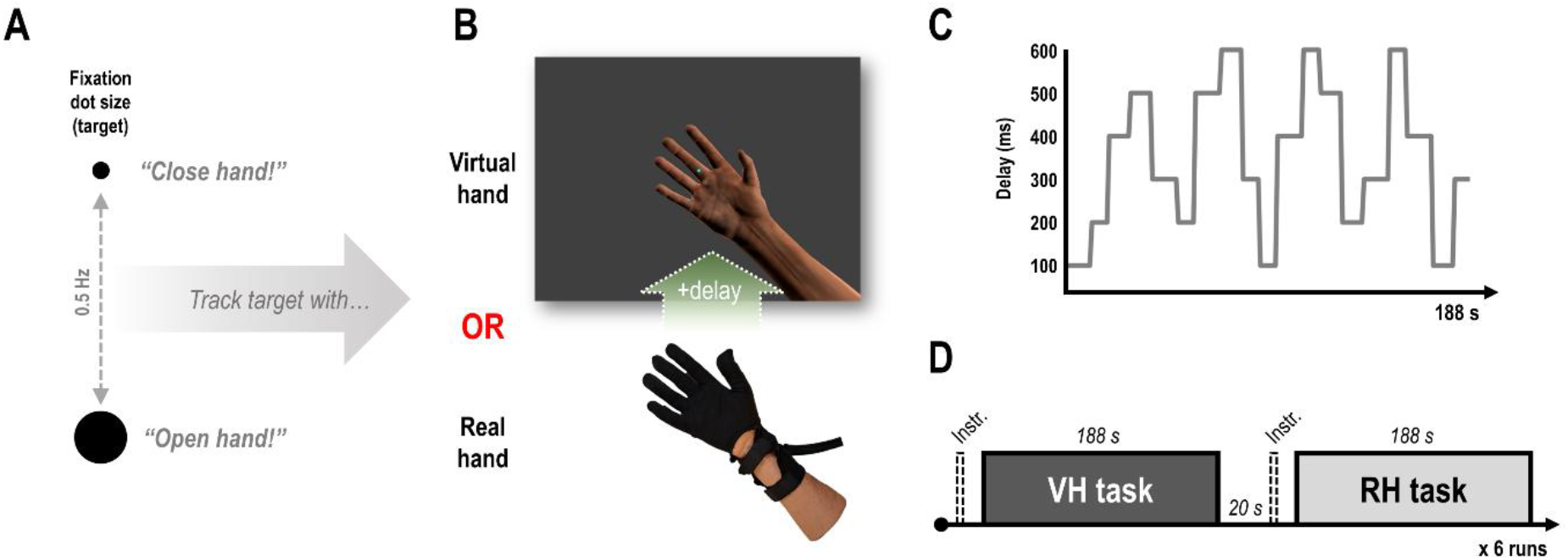
Experimental task and design. **A:** The participants had to match the phase of an oscillating dot with recurrent right-hand (open-and-close) movements. **B:** Via an MR compatible data glove on their right hand, participants controlled a photorealistic virtual hand model. The virtual hand movements were permanently delayed with respect to the real hand movements. Thus, only one of the hands (virtual or real) could be aligned with the target oscillation at a time, while the other one consequently moved out of phase. **C:** During the continuous movement task, the amount of visuomotor delay varied between 100 and 600 ms in a roving oddball fashion, requiring repeated adaptation in the VH task. **D:** Participants were instructed to match the target oscillation with the movements of either the virtual hand (VH) or their real, unseen hand (RH). Thus, participants either had to attend to, and try to adapt to the visuomotor delays (VH task), or to ignore them (RH task). Reprinted from Vigh and Limanowski (2025).

Here, we subjected these data to dynamic causal modeling (DCM, Friston, 2003) to test whether the observed task dependent regional BOLD signal changes could be explained in terms of changes in interregional communication in a network comprising these areas, potentially in line with the assumption of cerebellar (or parietal) forward predictions.

## Methods

### Participants

We re-analyzed the fMRI data acquired by Vigh and Limanowski (2025). Twenty healthy, right-handed participants (15 female, mean age = 28.1 years, range= 22-38, normal or corrected-to-normal vision) had taken part in the experiment; which had been approved by the ethics committee of the Technische Universität Dresden.

### Experimental task and design

Participants had to perform a hand-target phase matching task; i.e., they had to match the phase (0.5 Hz) of an oscillating fixation dot with right-hand grasping movements (Fig. 1A). Participants wore a data glove on their unseen hand, which transmitted finger flexion data to a virtual hand presented on screen. Thus, the participants could move the fingers of the virtual hand (Fig. 1B). During the continuous movement task (in blocks of 188 s, see below), the movements of the virtual hand were permanently delayed with respect to the actually executed hand movements; this implied a constant visuomotor and visuoproprioceptive mismatch. Crucially, participants were instructed to match the target oscillation with their unseen, real hand or with the seen virtual hand. Note that, due to the delay added to the visual movements, only one of the two hands (virtual or real) could be aligned with the oscillating target at a time— the other hand would consequently move out of phase with the target. For instance, to align the virtual hand with the target, participants had to try to adapt their real hand movements (phase shift them to compensate for the delay); to align the real hand, participants had to ignore the visual feedback delay. Thus, the task instruction determined whether or not visual feedback delays were task relevant, and whether or not participants needed to try to adapt to them or ignore them. The amount of visual feedback delay was repeatedly varied between 100-600 ms in a roving oddball fashion, in pseudorandomized sequences, requiring repeated visuomotor adaptation (in the VH task, Fig. 1C). Each of 6 delay sequences consisted of 94 movement cycles (188 s) and was used for the VH and RH task alike (Fig. 1D).

As expected, our previous results (Vigh & Limanowski, 2025) showed that participants significantly more strongly phase-shifted their real hand movements depending on the amount of delay in the VH > RH task. Furthermore, the group-level statistical parametric mapping (SPM) analysis of the fMRI data had identified several key regions that showed stronger activation and, partly, stronger delay related responses in the VH > RH task: Firstly, the left IPS and the bilateral middle and inferior occipital gyri showing a significantly (*p*_*FWE*_ <.05) stronger BOLD signal during the VH > RH task. This suggested an involvement of these regions in visuomotor integration and related attentional control during the VH > RH task. Here we assign these occipital activations the label “V5”, as they very likely correspond to these motion sensitive areas. Secondly, we identified regions in the left (lobule VIIIa) and right cerebellum (lobule VI/VIIa) showing an increased BOLD signal correlation with the parametric delay regressor in the VH > RH task; these regions also showed increased BOLD signal during the VH > RH task in a conjunction search (i.e., masking two contrasts with each other at *p* <.001, uncorrected). Thus, the cerebellar response increase was elevated during the VH > RH task in a delay-dependent manner, suggesting a process related specifically to delay compensation and visuomotor adaptation. No other regions showed significant effects of the VH > RH task or delay regressor.

### DCM analysis

The focus of our DCM connectivity analysis was on explaining the stronger activations and delay correlations observed during the VH > RH task in terms of network interactions; i.e., changes in connectivity among our regions of interest (ROIs). The above five ROIs, the left IPS, the bilateral V5, and the bilateral cerebellum, were selected as nodes for our DCM analysis (Fig. 2A). For each ROI, we extracted the BOLD signal time series, concatenating the 6 experimental runs, from 4 mm radius spheres centered on the individual peak voxel (i.e., the strongest participant specific effect) within 10 mm of the group cluster’s peak effect; i.e., for the contrast Task VH > RH for the IPS and V5s, and for the contrast Delay VH > Delay RH for the cerebella. The mean MNI coordinates of the peak effects with associated standard deviations were: left IPS (x = −33.0±4.8, y = −45.7±4.8, z = 48.2±4.9); left V5 (x = −45.0±4.4, y = −77.9±4.7, z = −3.1±4.6); right V5 (x = 42.0±5.0, y = −69.6±4.6, z = −14.2±4.9); left cerebellum (x = −37.8±3.8, y = −60.3±5.3, z = −26.4±4.2); right cerebellum (x = 37.1±4.9, y = −41.9±4.3, z = −31.7±5.2). To reduce noise, we extracted the time series only from voxels showing the desired effect at a statistical threshold of p <.05, uncorrected (cf. Vossel et al., 2012); in some cases, we had to increase the statistical threshold to reveal significant voxels (between p <.1 to p <.7; cf. Zeidman et al., 2019a). The time series were adjusted for effects of no interest (movement regressors and session means).

**Figure 2:**
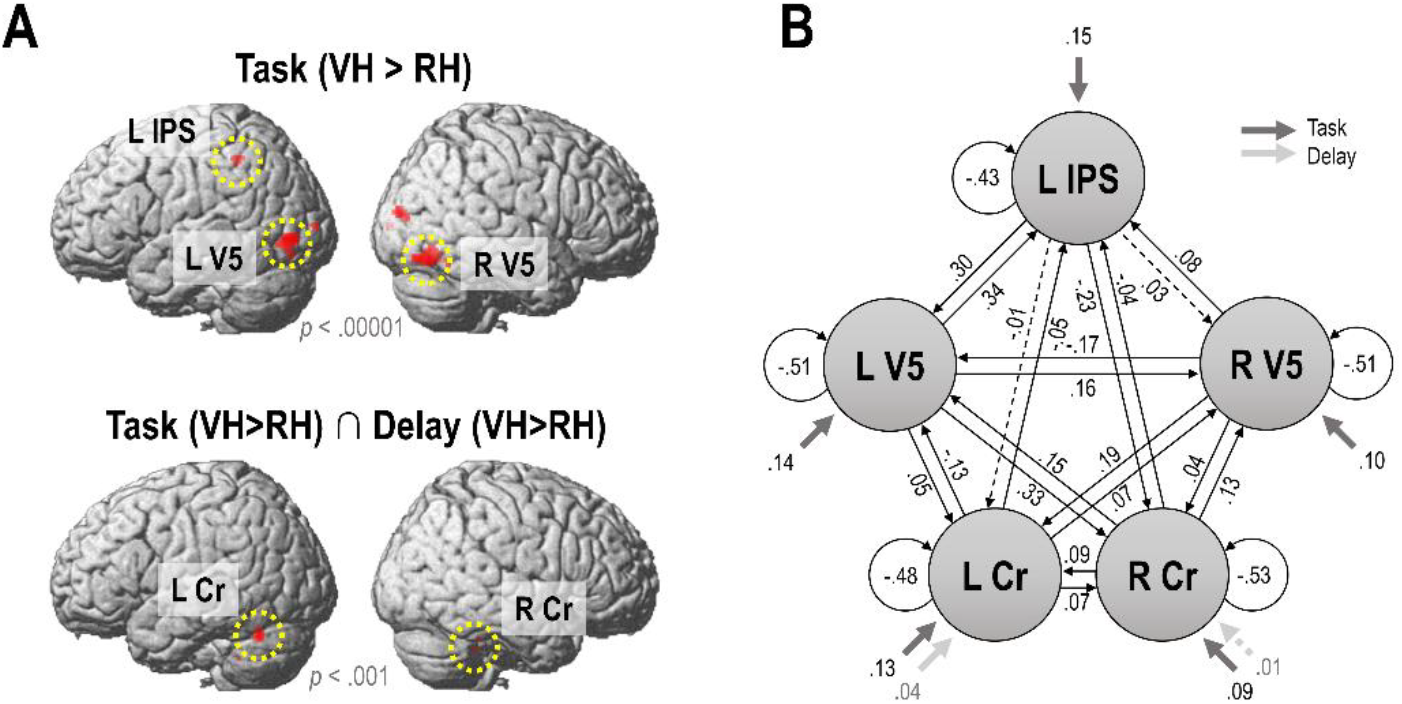
DCM architecture. **A:** Based on the significant effects obtained from the group-level mass univariate SPM analysis, five nodes of interest (circled in yellow) were selected for the DCM analysis: The left IPS and the bilateral V5 showed significantly increased activation during the VH > RH task (*p*_*FWE*_ <.05, render thresholded at *p* <.00001 for display purposes); the bilateral cerebellum (Cr) a stronger activation by delays in the VH > RH task and an increased activation by the VH > RH task per se (*p* <.001, uncorrected). **B:** DCM model architecture showing driving inputs (C matrix parameter estimates) and baseline network connectivity (A matrix parameter estimates) among the five nodes of the full model. Dashed lines indicate connections with only weak evidence (posterior probability <.75). This architecture was the basis for identifying task-dependent modulations of connection strengths (Fig. 3).

For the intrinsic connectivity of our model (A matrix), we allowed bilateral connections between all nodes of the network, and each node received an inhibitory ‘self’-connection. Driving inputs (C matrix) of the task (i.e., regressor consisting of the VH + RH task) were allowed to each region, as each of the ROIs showed activation during the task > rest in our initial GLM analysis. Driving inputs of the delay effect (i.e., regressor consisting of the parametric delay effect in the VH + RH task) were allowed to the bilateral cerebellum, which showed a main effect of delay in the GLM analysis. Figure 2B shows the intrinsic connectivity of the network at baseline, and the effects of the driving inputs.

In this model, we then defined two modulatory inputs (B_1_ and B_2_ matrices): of the VH and the RH task, respectively, which could act upon all between-region connections. The inputs were not mean centered, so the B matrix (modulatory) parameters can be interpreted as in-or decreases in connection strength, during the VH or RH task, relative to baseline activity captured by the A matrix. The full model was then estimated (inverted) for each participant; on average, these models explained 25.2% of the experimental variance. To determine the optimal explanation of our experimental effects (i.e., the combination of B matrix connectivity parameters best explaining the observed BOLD signal time differences), we used Bayesian model reduction (BMR; Friston et al., 2016) within a parametric empirical Bayesian (PEB; Zeidman et al., 2019a,b) approach, where a ‘full’ model is estimated for each participant and inference is subsequently performed on reduced (‘nested’) models. In the BMR, reduced models consist of various different combinations of B matrix parameters switched off; the automatic ‘greedy search’ algorithm iteratively discards those parameters that do not contribute to model evidence—i.e., each connection parameter receives a posterior probability—until discarding further parameters decreases model evidence (Zeidman et al., 2019b). In other words, a model with the retained parameters (modulated connections) explains the data (BOLD signals) better than a model without them. Then, we calculated a Bayesian model average (BMA) over the 256 models of the final iteration of the search; i.e., the retained parameter estimates reflect the averages from different models weighted by the models’ respective posterior probabilities (Penny et al., 2010). We thresholded the results of this BMA at a posterior probability of >.75 to highlight parameters with positive evidence for a contribution to model evidence; i.e., a model with this parameter would notably outperform a model without it. For completeness, we report all retained parameters in Table 1, including those with posterior probabilities <.75. Finally, to illustrate the strength of the notable cerebellar connectivity differences between the VH and RH task, we computed Bayesian contrasts.

**Table 1.**
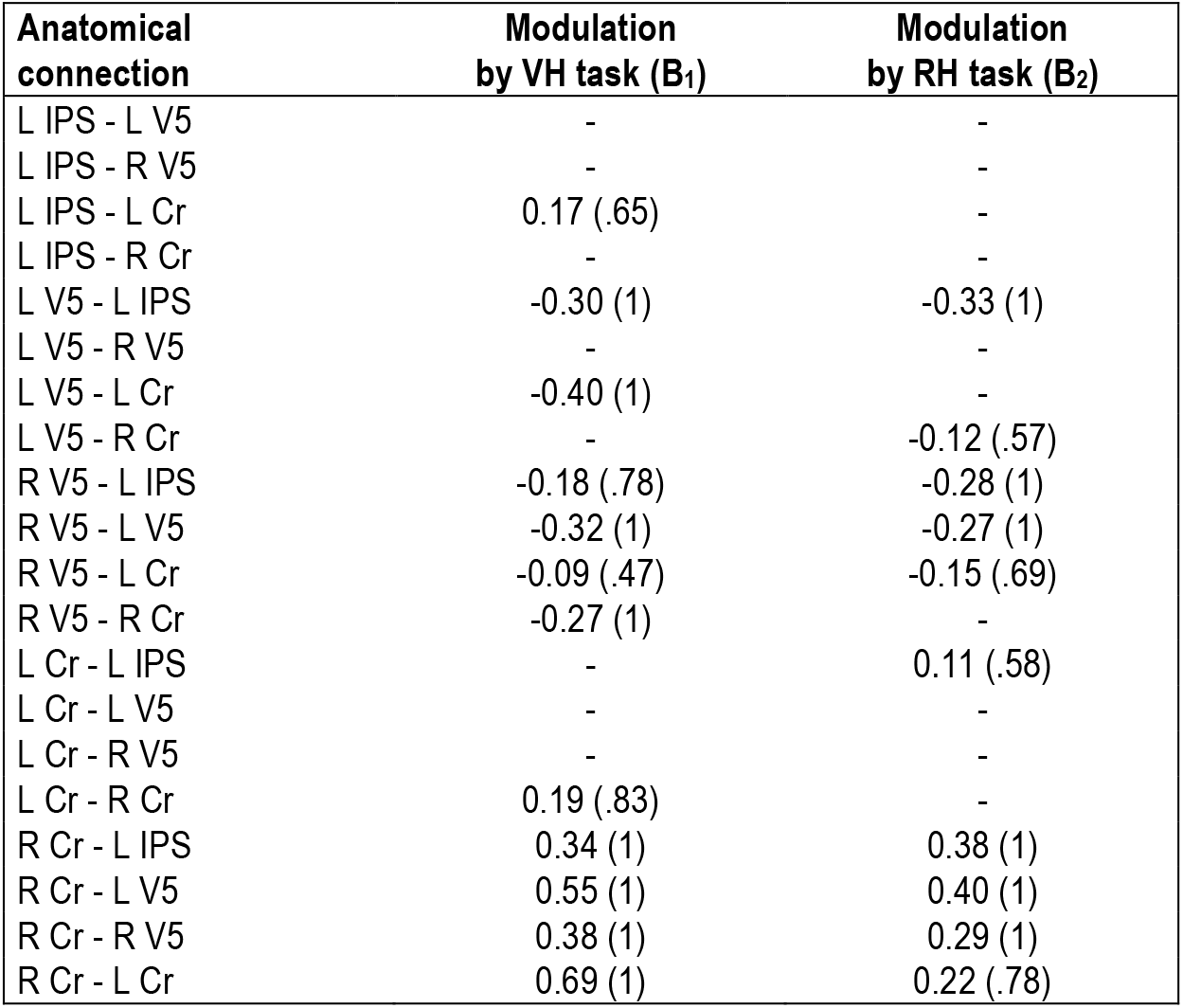
Bayesian model averages of parameters retained in the BMR with associated posterior probabilities in brackets (unthresholded).

## Results

BMR (Fig. 3A) retained several modulatory parameters, i.e., in- or decreases in connectivity strength that contributed to model evidence, in each task: Both tasks were associated with a positive influence of the right cerebellum on all other nodes of the network, with an inhibitory influence of the bilateral V5 on the IPS, and of the right of the left V5.

**Figure 3:**
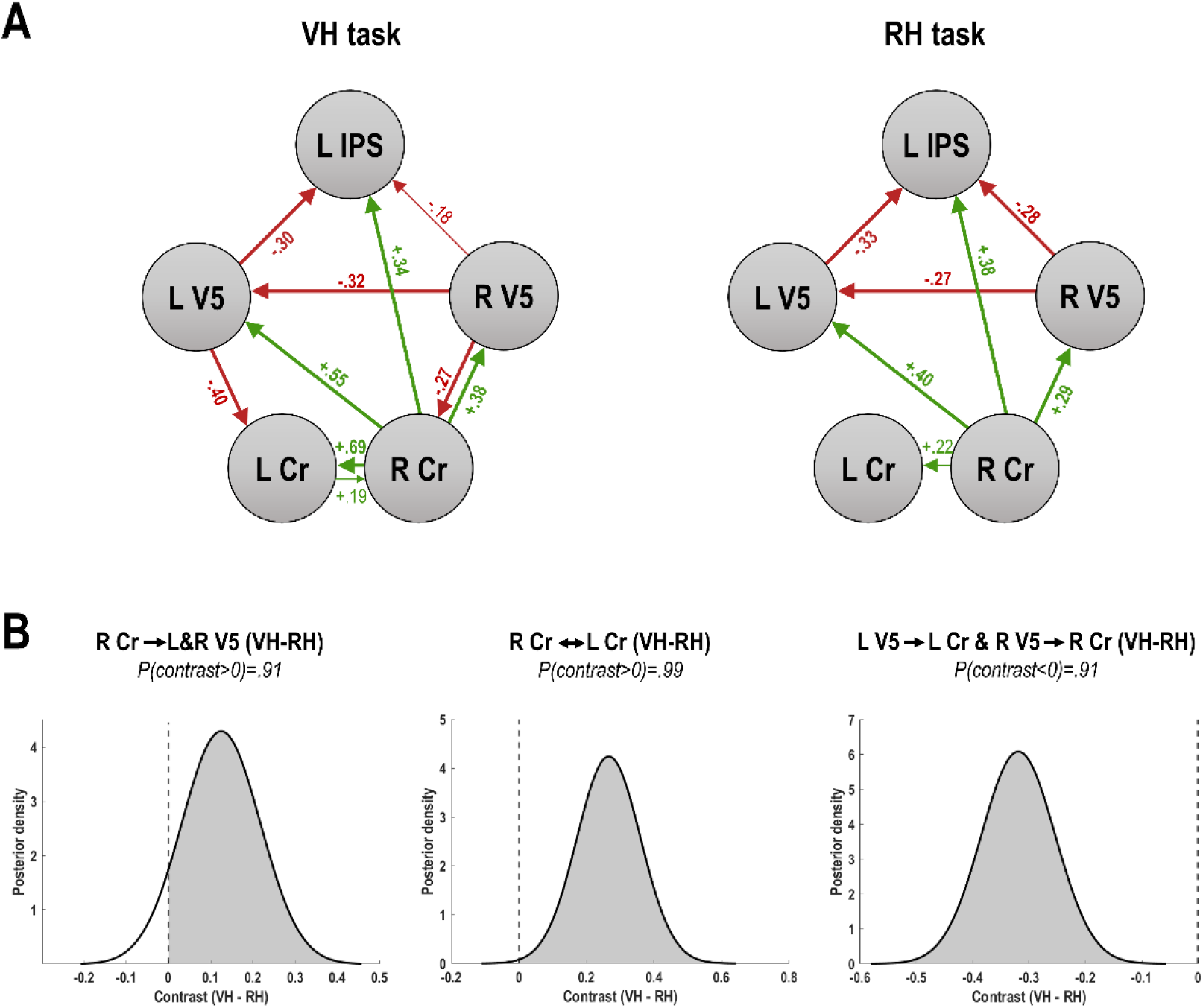
Task dependent connectivity changes. **A:** Results of the automated search over reduced models, showing the retained connections of the winning model. Green and red colors reflect positive and negative modulations, respectively; thick lines indicate estimates with a posterior probability >.95, thin lines those with a posterior probability of >.75. The parameter estimates are Bayesian model averages reflecting the in- or decreases from baseline connectivity. See Table 1 for details. **B:** Bayesian contrasts over parameter estimates (VH task - RH task) showed strong evidence for that the connections from the right cerebellum to the bilateral V5 (left plot), and the mutual connections between the left and right cerebellum (middle plot) were more excitatory in the VH task compared with the RH task (indicated by positive values of the posterior means); while the connectivity from each V5 to the ipsilateral cerebellum was more inhibitory (right plot, indicated by negative values).

Crucially, three kinds of connections showed marked differences between tasks: Firstly, the mutual excitatory influence among the right Cr and the left Cr was more strongly increased (from a slightly positive baseline, Fig. 2B) in the VH>RH task. Likewise, secondly, the excitatory influence of the right Cr onto both V5s was more strongly increased in the VH>RH task. Thirdly, in each hemisphere, the V5 exhibited an inhibitory influence on the Cr in the VH, but not the RH task (where these connections were removed i.e. set to 0 by the BMR). Bayesian contrasts over joint parameters implementing the above three key task dependent differences in (cortico-)cerebellar connectivity showed that they had high posterior probabilities of being non-zero (Fig. 2B). See Table 1 for an overview of all parameter estimates with associated posterior probabilities.

## Discussion

Our results highlight the role of network interactions of the (right) cerebellum in a task involving adaptation to delayed visual movement feedback (VH task), compared with moving while ignoring those delays (RH task). The right cerebellum had an increased excitatory effect on all other nodes of the network in both tasks, in line with the idea that cerebellar-to-cortical communication is crucial in on-line action control (see Introduction). Specifically, we propose the excitation suggests that cerebellar state estimates or (sensory) predictions drive activity in the IPS and in the motion-sensitive V5 in a ‘top-down’ fashion (cf. Kellermann et al., 2012).

More importantly, several of these connections differed between tasks. Thus, the increased excitatory effect of the right cerebellum on the bilateral V5 in the VH>RH task indicated a selectively increased sensitivity of the motion sensitive V5s to cerebellar predictions. Furthermore, there was an increased mutual excitation among the cerebellar hemispheres in the VH>RH task. As the driving effect of delays was estimated to predominantly enter the left cerebellum, this interaction could indicate enhanced delay processing (cf. Leube et al., 2003; Arikan et al., 2019) across the cerebellar hemispheres in the visuomotor adaptation task. In line with this interpretation, increased connectivity between the left and right cerebellum has been reported in DCM studies during social action prediction (Haihambo et al., 2025; see also Metoki et al., 2022), which involves similar (visual) predictive processes. Interactions between cerebellar hemispheres could also be related to other processes such as timing prediction and sensorimotor synchronization (cf. O’Reilly et al., 2010, Manto et al., 2012; Pollock et al., 2006; Sokolov et al., 2017). Together, the increased excitatory influences of the (right) cerebellum on other nodes of the visuomotor control hierarchy during the visuomotor adaptation (vs no-adaptation) task support the hypothesis that it is predicting dynamic sensory movement consequences (here, of visual motion) for adaptive, visually guided on-line control.

Conversely, the V5s inhibited the cerebellum during the VH, but not the RH task. The V5 also had inhibitory effects on the IPS, in both tasks. An increased inhibition of the PPC by higher-level visual regions has been observed in other DCM studies, e.g., during attention to motion or visual perception (Kellermann et al., 2012; Dijkstra et al., 2017). Kellermann et al. (2012) proposed an interpretation in terms of a reduced parietal sensitivity to bottom-up prediction errors (also from the V5 in their case) during attention to motion. A similar interpretation could be applied to our results; i.e., the higher-level cerebellum’s predictive models would be influenced less by signals about (delayed) visual motion in the VH than the RH task, because they were adapting to the delays. However, the parameter estimates relative to baseline suggested an actual inhibitory effect of the V5 on the cerebellum, not just a reduction in excitation. Thus, an alternative potential explanation could be that input from the V5 inhibits cerebellar activity devoted to forward modeling, potentially to accommodate other information or processes. This would, tentatively, fit with DCM findings of an inhibition of the cerebellum by the precuneus during the observation of (social) actions consistent>inconsistent with expectations (Li et al., 2025). Disentangling between these interpretations requires further studies with complementary experimental designs.

It should be noted that, as customary, we applied DCM to regions showing experimental variance in their BOLD signal, thus neglecting other parts of the visual and visuomotor hierarchy. Thus, our results should be interpreted together with previous work showing the importance of cerebellar connectivity with premotor regions during visuomotor (de)adaptation (Tzvi et al., 2020). We found no BOLD signal changes in somatosensory regions, but other studies suggest changes in brain connectivity underlying the relative weighting of vision vs somatosensation during intersensory conflicts (Limanowski et al., 2015; Limanowski & Friston, 2020; Pamplona et al., 2025). Finally, studies using perceptual tasks have revealed the effects of prediction and attention on connectivity within earlier parts of the visual hierarchy (Kok et al., 2012; Kellermann et al., 2017; cf. Büchel et al., 1998), in the dorsal or ventral streams (Vossel et al., 2012; 2015; Zbären et al., 2024), or the motor system (Bencivenga et al., 2021). Our results add to these previous findings evidence for that cerebellar signals to visual (and sensorimotor parietal) regions convey predictive signals during visuomotor adaptation.

## Acknowledgments

We thank Peng Wang and Eric Bendt for assistance with the data analysis.

